# Standing wave mesoscopy

**DOI:** 10.1101/2023.03.08.531677

**Authors:** Shannan Foylan, Jana Katharina Schniete, Lisa Sophie Kölln, John Dempster, Carsten Gram Hansen, Michael Shaw, Trevor John Bushell, Gail McConnell

## Abstract

Standing wave (SW) microscopy is a method that uses an interference pattern to excite fluorescence from labelled cellular structures and produces high-resolution images of three-dimensional objects in a two-dimensional dataset. SW microscopy is performed with high magnification, high numerical aperture objective lenses, and while this results in high resolution images, the field of view is very small. Here we report upscaling of this interference imaging method from the microscale to the mesoscale using the Mesolens, which has the unusual combination of a low magnification and high numerical aperture. With this method, we produce SW images within a field of view of 4.4 mm x 3.0 mm that can readily accommodate over 16,000 cells in a single dataset. We demonstrate the method using both single-wavelength excitation and the multi-wavelength SW method TartanSW. We show application of the method for imaging of fixed and living cells specimens, with the first application of SW imaging to study cells under flow conditions.

## 1. INTRODUCTION

Standing wave (SW) microscopy uses axially structured illumination to produce a high-resolution topographical map of cellular structures. ^1^ The excitation field is generated by two beams from an individual or different optical sources which interfere and form a SW, creating a sinusoidal excitation pattern in the axial direction. Only fluorescent molecules that coincide with the anti-nodal plane maxima of the SW will be excited and emit fluorescence. ^1^

Equidistant planes of fluorescence emission are generated within the fluorescently-labelled specimen that are perpendicular to the optical axis and are separated by

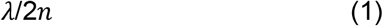

where λ is the wavelength of excitation light and *n* is the refractive index of the medium in which the light is propagating. The individual anti-nodal planes are

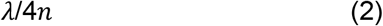

thick at the full-width at half maximum (FWHM). This FWHM value of anti-nodal plane thickness is commonly used as the definition of the axial resolution in SW imaging. ^1^

SW microscopy has been used previously to study structures including cell topography and the actin cytoskeleton. ^1,2^ Both single-wavelength and, more recently, multi-wavelength SW microscopy (TartanSW) have been reported, with axial sampling densities of 50% to 98% respectively. ^3^ As with conventional cell imaging, high magnification, high numerical aperture lenses are normally used for SW microscopy. Such lenses provide a field of view (FOV) on the order of a few tens of μm in diameter, which limits the number of cells that can be observed in a single image. As described above, the axial super-resolution in SW microscopy is independent of numerical aperture (NA), but the lateral resolution, *r*, of the widefield image follows the standard Rayleigh resolution criterion ^4^

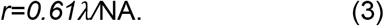

Ideally, an imaging lens with a high numerical aperture and a low magnification is required for SW microscopy to visualise large cell populations. Large data sets are needed to accurately visualise and analyse cell population heterogeneity and detect rare abnormalities in large cell populations. ^5^

The Mesolens ^6^ is a complex objective with a magnification of 4x and an NA of 0.47. This gives a large improvement in image information content compared with commercial objective lenses. The Mesolens is flat field corrected over a 5.5 mm FOV, and is chromatically corrected across the visible spectral range. We have previously reported the Mesolens as the imaging objective in confocal microscopy, ^6,7^ but it is time-consuming to acquire three-dimensional image data with point-scanning illumination. For example, confocal imaging with the Mesolens within a reduced FOV to study a 4.4 mm x 3 mm x 20 μm thick specimen at Nyquist sampling (4 pixels/μm in xy, and 10 images in z) with a pixel dwell time of 1 μs and a frame average of two takes almost twelve hours. ^6^ The time for acquisition of this volume with commercial setups that employ stitching and tiling methods at the same resolution is similarly long. Light-sheet mesoscopy has reduced the overall acquisition time considerably, although this requires a complex optical arrangement and thus far has only been demonstrated for a single excitation wavelength. ^8^ Multi-colour, two-dimensional imaging with the Mesolens has been demonstrated with a sensor-shifting camera, producing Nyquist sampled images in under 15 seconds. ^9^ Colour brightfield and widefield epi-fluorescence mesoscale imaging of blood films and large biofilms have been reported but, as with conventional microscopy, these methods are best suited for optical imaging of thin specimens, and they provide only two-dimensional information about the cellular structure. ^10,11^

Here, we show how combining the Mesolens with the SW illumination method enables high-resolution mesoscale SW imaging of large cell populations in both fixed and live specimens. We report multi-wavelength SW imaging of fixed fibroblast cells, and single-wavelength SW imaging of more than 16,000 red blood cells in a single image that was acquired in under 15 seconds. Such cells continuously experience multiple types of mechanical forces including shear stress in the circulatory system, and these forces can influence regeneration. ^12^ We therefore also explore the application of SW mesoscopy to monitor cell behaviour under flow conditions. Using the SW method, we show that it is possible to obtain multi-plane illumination at the mesoscale, and hence visualise the three-dimensional contours of cell structures from large numbers of cells in a single dataset.

## 2. METHOD

### 2.1 Non-biological test specimen preparation

An uncoated silica plano-convex lens with a focal length of f=400 mm (400 PQ 25, Comar Optics, Haverhill, UK) was prepared as described previously, ^2^ using a 10 μM solution of DiO (3,3’-Dioctadecyloxacarbocyanine Perchlorate) (D275, Thermo Fisher, Paisley, UK) instead of Atto 532 NHS-ester. The lens was placed with the curved surface in contact with a first surface reflector (TFA-20C03-10, Laser 2000, Huntingdon, UK). We chose this mirror for its high flatness (λ/10) and the absence of a protective coating, which we previously found to be problematic for SW microscopy. ^3^ This specimen was imaged with no fluid mountant, i.e. in air, to ensure that the lens was positioned with the planar surface orthogonal to the optical axis of the Mesolens.

### 2.2 Biological specimen preparation

#### 2.2.1 Fibroblast cells

Fibroblast cells (3T3-L1, CL-173) were grown in vented capped tissue culture flasks containing DMEM (Gibco 10567-014, Thermo Fisher, Paisley, UK) supplemented with 10% FBS (Labtech, Heathfield, UK) and 1% Penicillin Streptomycin (Gibco, Thermo Fisher, Paisley, UK) and 1% L-glutamine (Thermo Fisher, Paisley, UK), and were incubated at a temperature of 37 °C in a 5% CO_2_ humidified cell incubator (Heracell VIOS CO_2_, Thermo Fisher, Paisley, UK).

The same type of mirrors used to image the lens specimen were prepared with a 1:500 dilution of fibronectin bovine plasma (F1171-2MG, Sigma Aldrich, UK). Mirrors were placed in 6 well plates with the reflective surface facing the open side of the well. Cells were seeded at the desired density and incubated for 24 h at 37 °C/ 5% CO_2_ to promote adherence to the coated mirror and then rinsed once with an appropriate buffer before staining.

For staining with fluorescein phalloidin (F432, Thermo Fisher, Paisley, UK), cells were rinsed twice with PBS (1× PBS, 5 min) and then fixed in 4% PFA (Paraformaldehyde, Sigma Aldrich, Dorset, UK) for 10 min. This was followed by rinsing three times in PBS (1× PBS, 5 min), and then blocking with 1% BSA in PBS for 30 min. Cells were then incubated for 20 mins under light-tight conditions in fluorescein phalloidin diluted 1:2,000 in 1% BSA/PBS buffer before being washed twice (1× PBS, 30 s) and stored in PBS at 4 °C until imaged. Prior to imaging, a 70 mm x 70 mm type 1.5 coverslip (0107999098, Marienfeld, Lauda-Koenigshofen, Germany) was placed on top of the cell specimen. This unusually large coverslip was required to support a 25 mm diameter layer of water that extended from the coverslip to the front element of the Mesolens. At a height of approximately 3 mm this layer of water was slightly thinner than the 3.1 mm working distance of the Mesolens, and was held in place with surface tension.

#### 2.2.2 Live red blood cells

Human erythrocytes were collected through a needle puncture from the fingertip, yielding around 0.5 ml blood, with ethical permission (NHS Biorep 548). A working solution of 5 μg/ml of FM 4-64 (T13320, Thermo Fisher, Paisley, UK) in water was prepared and kept at room temperature. An Eppendorf tube containing the red blood cells sample was kept at room temperature, and 100 μl of the staining solution was added. After 1 minute, either 10 μl or 50 μl of the stained cell suspension was added directly to the same mirrors used to image the lens specimen and fibroblasts. The low-volume cell suspension was used to create a specimen in which cells were almost all in contact with the mirror, while the high-volume suspension was used to image cells in a flow environment. This was required because, to our knowledge, there are no flow chambers of a size suitable for imaging with the Mesolens that incorporate the first surface reflector needed for SW imaging. The mirror prepared with the low-volume cell suspension was allowed to settle for 2 minutes at room temperature, while the specimen prepared with the high-volume cell suspension was imaged immediately, with no time for the cells to settle on the mirror. In both cases a 70 mm x 70 mm type 1.5 coverslip (0107999098, Marienfeld, Lauda-Koenigshofen, Germany) was placed on top of the cell specimen.

### 2.3 Mesoscale imaging

A diagram of the SW mesoscopy imaging setup is shown in Figure 1. The Mesolens was used in widefield epi-illumination mode with a high-brightness multi-wavelength light emitting diode (LED) illuminator (pE-4000, CoolLED, Andover, UK) to sequentially deliver light with wavelengths of 385 ± 15 nm, 430 ± 20 nm, and 490 ± 20 nm to the specimen plane. The resulting fluorescent emission from a specimen was propagated through multi-bandpass (Pinkel-type) ^13^ chromatic reflector and barrier filters. The Pinkel design was used so that fluorescence of different excitation and emission spectral ranges could be recorded sequentially without the image movement produced by conventional replaceable filters. We used these to detect fluorescence emission at 417 ± 10 nm, 460 nm ± 10 nm, 525 nm ± 25 nm, and 635 ± 20 nm. ^13^ As we have shown previously, it is unnecessary to use a light source with a long coherence length for SW microscopy, and light emitting diodes are suitable for this purpose. ^14^

**Figure 1.**
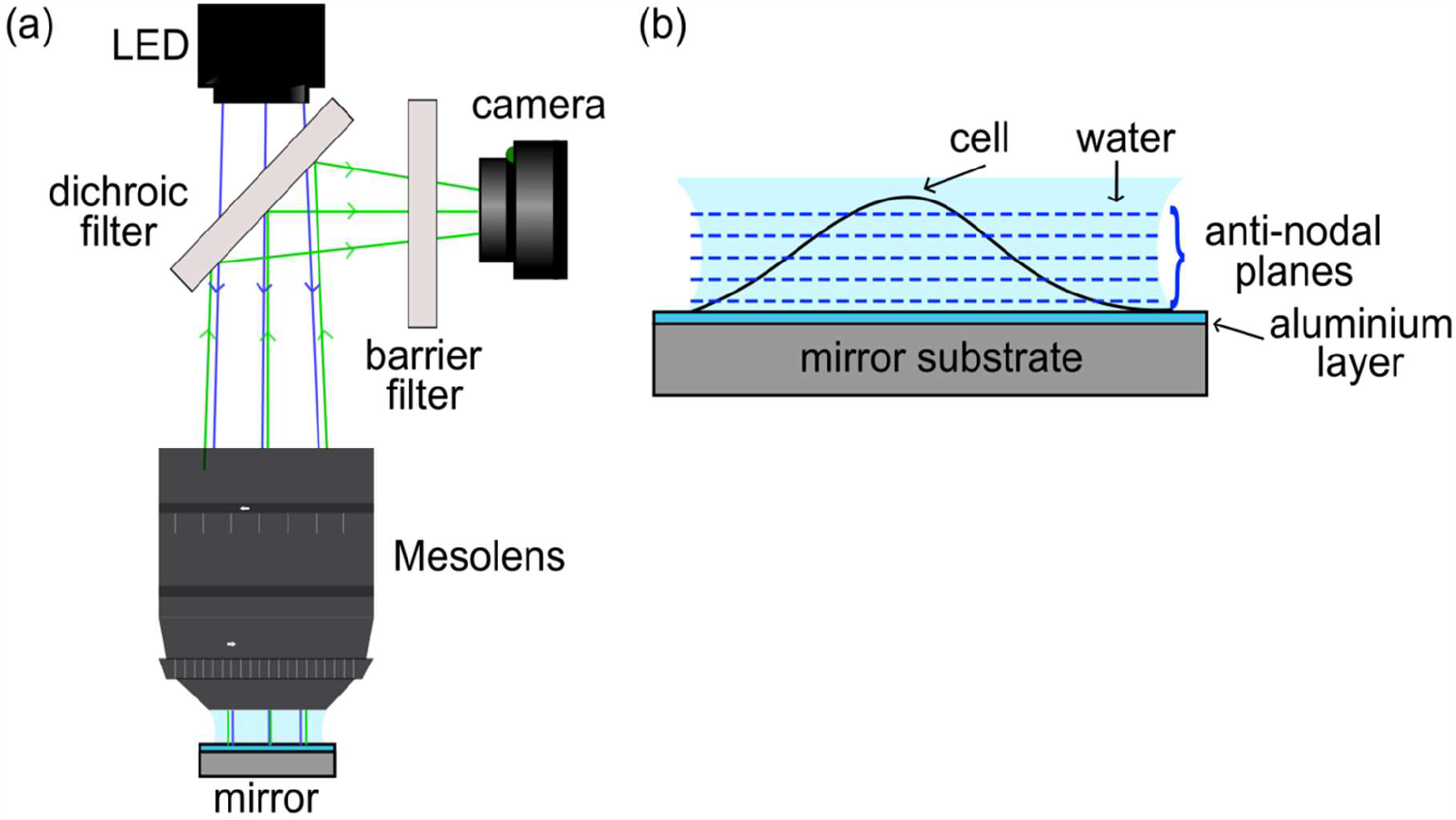
Experimental setup for standing wave mesoscopy. (a) The ray diagram for the LED illumination is shown in blue, with the ray diagram showing fluorescence emission in green. (b) Schematic view of a mammalian cell specimen attached to the aluminised surface of the mirror showing the anti-nodal planes of high intensity (dotted lines). The spacing of the anti-nodal planes is not shown to scale.

Single-wavelength SW images of the lens specimen were acquired with 490 ± 20 nm excitation with 18.3 mW average power at the specimen plane, with a 1,000 ms camera exposure and gain of 50x. TartanSW images of the fibroblast cells were obtained using sequential wavelength excitation from all three LEDs: 385 ± 15 nm with an average irradiance of 1.14 kW/m^2^, 430 ± 20 nm with an average irradiance of 234 W/m^2^, and 490 ± 20 nm with an average irradiance of 371 W/m^2^, with all power values measured at the specimen plane. The exposure time was longest for the shortest wavelength at 5,000 ms, and this was decreased to 800 ms at 430 ± 20 nm, and 200 ms for the longest wavelength of illumination. The camera gain was set to 80x for imaging at 385 nm, and this was decreased to 40x for imaging with both 430 ± 20 nm and 490 ± 20 nm. Images from the three different channels were saved individually for analysis and processing. Images of red blood cell specimens were acquired with 490 ± 20 nm excitation at 20% LED power from the illuminator with a 100 ms camera exposure, and a gain of 40x. Time-lapse recording of the red blood cell specimens was also performed, with 10 images acquired from the high-volume cell suspension preparations, and 30 images acquired from the low-volume cell suspensions using a 30 s time interval between recordings.

To capture the large, high-resolution images produced by the Mesolens we used a chip-shifting camera sensor (VNP-29MC; Vieworks, Anyang, Republic of Korea) which records images by shifting a 29-megapixel CCD chip in a 3 × 3 array. ^9^ In this mode, the sampling rate was 4.46 pixels/μm, corresponding to a 224 nm pixel size, satisfying the Nyquist sampling criterion. The minimum frame time of the Vieworks camera was 200 ms resulting in an acquisition time for one full FOV image with 9x pixel shift of 1,800 ms, excluding the time needed to transfer the image data from the camera to the PC which usually took on the order of 10 s. Therefore, in practice, the acquisition of one image took 12–15 s. For three-colour imaging of the fibroblasts, false-colour merging of each channel (260 Megapixels, 506 MB for each image) took approximately 5 s.

The fibroblast cell specimens were mounted in PBS (n = 1.34) for imaging. All specimens were imaged with water immersion, with the correction collars of the Mesolens set to minimise spherical aberration.

### 2.4 Image processing

The SW images of the plano-convex lens specimen were contrast adjusted using the Contrast Limited Adaptive Histogram Equalization (CLAHE) ^15^ function in FIJI ^16^ with the default parameters (blocksize = 127, histogram bins = 256, maximum slope = 3.00). For further analysis in MATLAB, images were cropped, centered on the zeroth order standing wave anti-node at the apex of the convex lens surface, and saved for further processing. The sequentially acquired images of each specimen for the different excitation wavelengths were contrast adjusted using the ‘Auto’ Brightness/Contrast function in FIJI. ^16^ To perform TartanSW mesoscopy, images were merged using the pseudo colours red, green and blue, with false colour red being used for the longest excitation wavelength and false colour blue for the shortest excitation wavelength of the imaging experiment.

To correct for inhomogeneous illumination across the FOV caused by the beam profile of the LED, each image was processed with a flatfield correction using SciPy. ^17^ Here, the background was assumed to be its Gaussian filtered image (σ=200) and the image was divided by this background image. A normalisation was performed to the 5th and 99.8th percentile of the image, including clipping of all pixel intensity values outside of the range of 0 to 1.

### 2.5 Image analysis

#### 2.5.1 Lens specimen

The SW images of the plano-convex lens specimens were analysed to confirm whether anti-nodal FWHM thicknesses at the mesoscale were comparable to values expected from theory. A previously published MATLAB script was used for analysis. ^14^ This code extracts the average line profile between the centre of the lens and the edge of the image and performs a radial average of fluorescence intensity values using the custom function *radialavg*. ^14^ In this work, 6,432 line profiles were measured. The average lateral intensity profile was then used to compute the axial intensity profile using Pythagoras’s theorem and the known geometry of the lens specimen to derive the equation

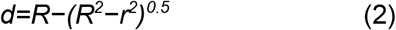

where *R* is the radius of curvature of the lens, *r* is the radial distance of each pixel from the centre of the image, and *d* is the axial distance from the mirror. ^14^ To determine the peak fluorescence intensity values for each anti-nodal plane we used the *findpeaks* function in MATLAB. ^14^ The average FWHM values were determined using the same *findpeaks* function then outputting and averaging the width argument from the function.

#### 2.5.2 Red blood cells

The number of red blood cells in the imaged FOV was measured by performing a manual threshold analysis in FIJI to generate a segmentation mask, and using the *Analyze Particles* function in FIJI ^16^ on this mask, objects with a diameter of between 4-8 μm were counted.

## 3. RESULTS

With the thin spherical fluorescent surface, it proved feasible to test the basic proposition that the Mesolens is compatible with single-wavelength-excitation SW mesoscopy. Figure 2 shows a single-wavelength-excitation SW image of the lens specimen obtained under 490 ± 20 nm excitation and simultaneous dual-band fluorescence detection at both 525 ± 25 nm and 635 ± 20 nm wavelength ranges using the Pinkel-type filter arrangement described in Section 2.4. Rings of fluorescent emission radiating from the centre of the FOV are clearly visible over the central 3.0 mm x 3.0 mm region of the image. The SW is likely to extend over the full FOV, but the curvature of the lens means that the anti-nodal planes are too close together in the lateral direction to be resolved at the edges of the FOV.

**Figure 2.**
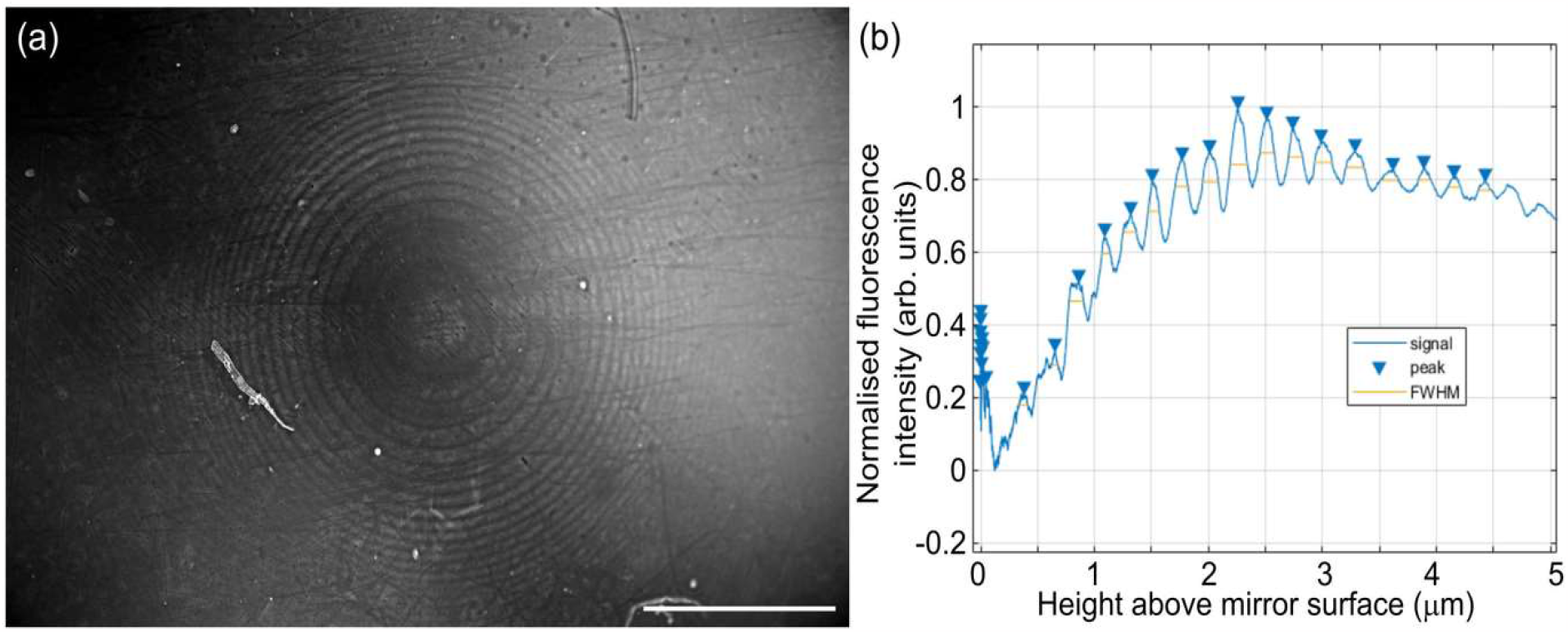
(a) Mesoscale SW image of the spherical surface of a plano-convex lens of focal length f=400 mm, with the curved surface stained with the fluorescent dye DiO. The curved surface of the lens was placed in contact with a first surface reflector and imaged in air using SW excitation. Fluorescence was excited at 490 ± 20 nm and detected between both 525 ± 25 nm and 635 ± 20 nm. Rings of fluorescent emission indicative of excitation with a SW are visible across the central 3.0 mm x 3.0 mm FOV of the image. Scale bar = 1 mm. (b) Fluorescence intensity plotted versus height from the mirror surface, showing evenly spaced peaks where the dye coincides with the anti-nodes of the illuminating SW pattern. The measured average anti-nodal FWHM thickness is 143 ± 19 nm.

The average FWHM thickness of the anti-nodal fluorescent rings was measured in the image to be 143 ± 19 nm (see Section 2.5). DiO has a broad emission spectrum with a peak at 506 nm. ^18^ Consequently, with the described setup, the image shown in Fig 2A visualises fluorescence from a broad spectral range. Using Equation (2) with n=1, FWHM thicknesses between 128 nm (for 512.5nm, the shortest detected emission wavelength) and 161 nm (for 645 nm, the longest detected emission wavelength) were expected according to theory. These predicted values are in good agreement with our experimental results.

Next, we applied SW mesoscale imaging to biological specimens. Figure 3(a) shows a full FOV pseudo-coloured image containing over 100 fibroblasts acquired using the TartanSW method obtained with sequential imaging and detection with the Mesolens after processing as described in Section 2.4. The image was downsampled to a size of 1000 pixels by 545 pixels for the purpose of presentation. Figures 3(b)-(d) show three full-resolution, zoomed-in regions of interest (ROIs) within the specimen. The TartanSW process results in multi-colour bands dependent on the axial distance of the fluorophore from the mirror surface which are visible across the full FOV. These data confirm suitability of the method at the mesoscale and for applications in imaging of large cell populations. While the multi-coloured anti-nodal planes are visible across the full FOV, we note that the contrast is slightly reduced in the top left of the image field. This is likely due to the specimen not being exactly orthogonal to the optical axis of the Mesolens. There is also disproportionate blue signal, resulting from 385 ± 15 nm illumination, in the bottom left of 3(a). This may be due to residue on the mirror surface. The cell on the left-hand side of Figure 3(d) has projections with dark regions in the centre of the filopodia, indicated by cyan arrows on the left side of the image. These are likely to be focal adhesions in direct contact with the mirror surface. ^19,20^

**Figure 3.**
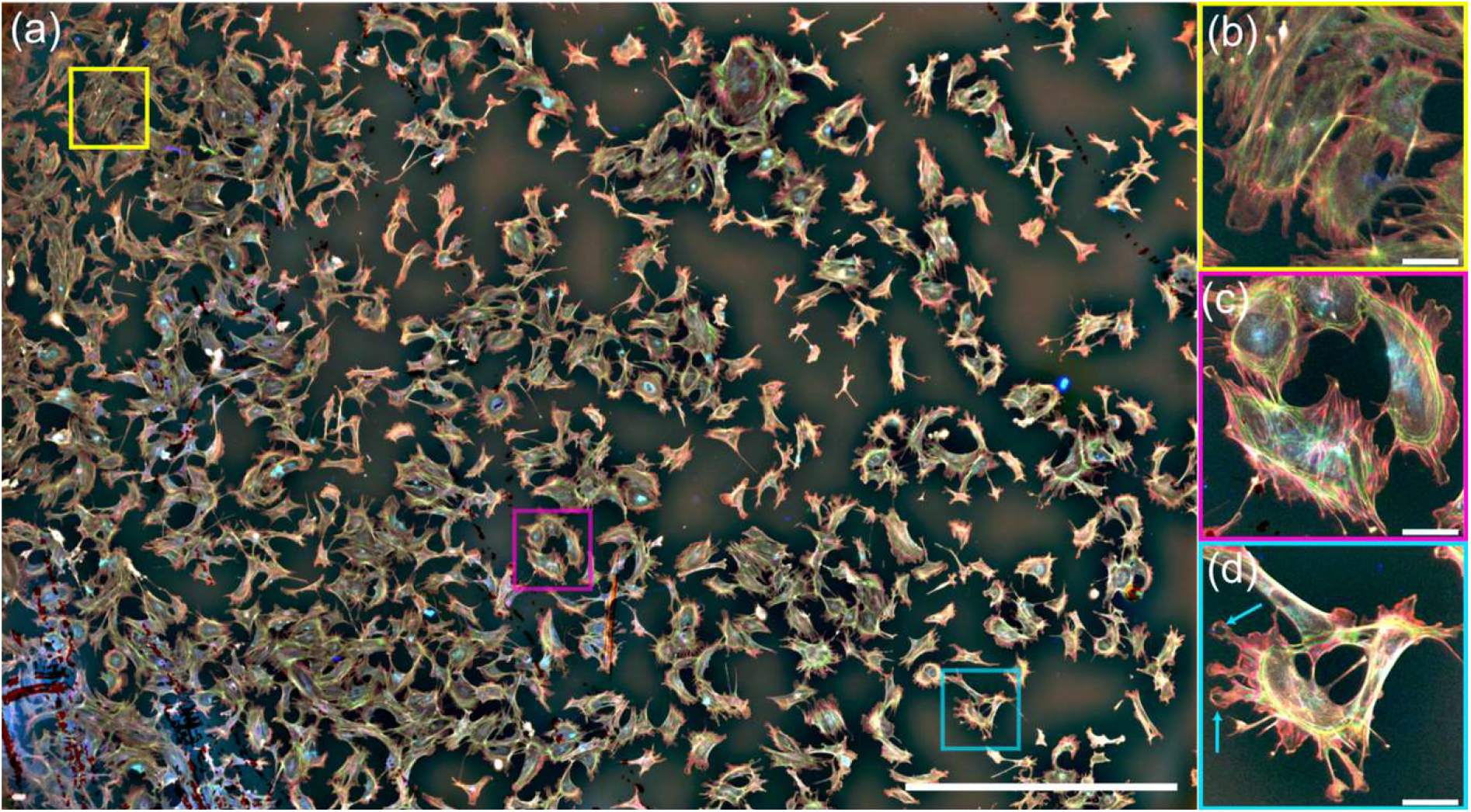
(a) Mesoscale TartanSW image of 3T3 cells stained with fluorescein phalloidin. Three ROIs are shown with a yellow box at the top left of the image, a magenta box close to the centre of the field, and a cyan box towards the bottom right of the field. Scale bar = 1 mm. (b-d) Digital zoom of the ROIs shown with the yellow, magenta, and cyan boxes respectively, with cyan arrows in (d) indicating possible focal adhesions. Scale bars for (b-d)= 50 μm.

Figure 4(a) shows a single-wavelength-excitation SW image of red cells obtained with the Mesolens, from a specimen prepared with the low-volume cell suspension (see Section 2.2.2). The large FOV relative to the small diameter of the cells means that it is difficult to see any cellular detail at this level of display zoom. Three ROIs are indicated by a yellow box in the top left of the image, a magenta box closer to the centre of the field, and a cyan box near the bottom right of the field. Figures 4(b)-4(d) show these digitally zoomed regions in more detail, with SW illumination visualizing distinct planes across the fluorescently-stained red blood cells. This confirms that our imaging method can be applied to small cells *in vitro* within an extended FOV. By conducting image segmentation as described in Section 2.5, we measure that there are 16,636 red cells in this single image. To our knowledge, this is the largest cell population reported thus far in a single dataset imaged with SW illumination.

**Figure 4.**
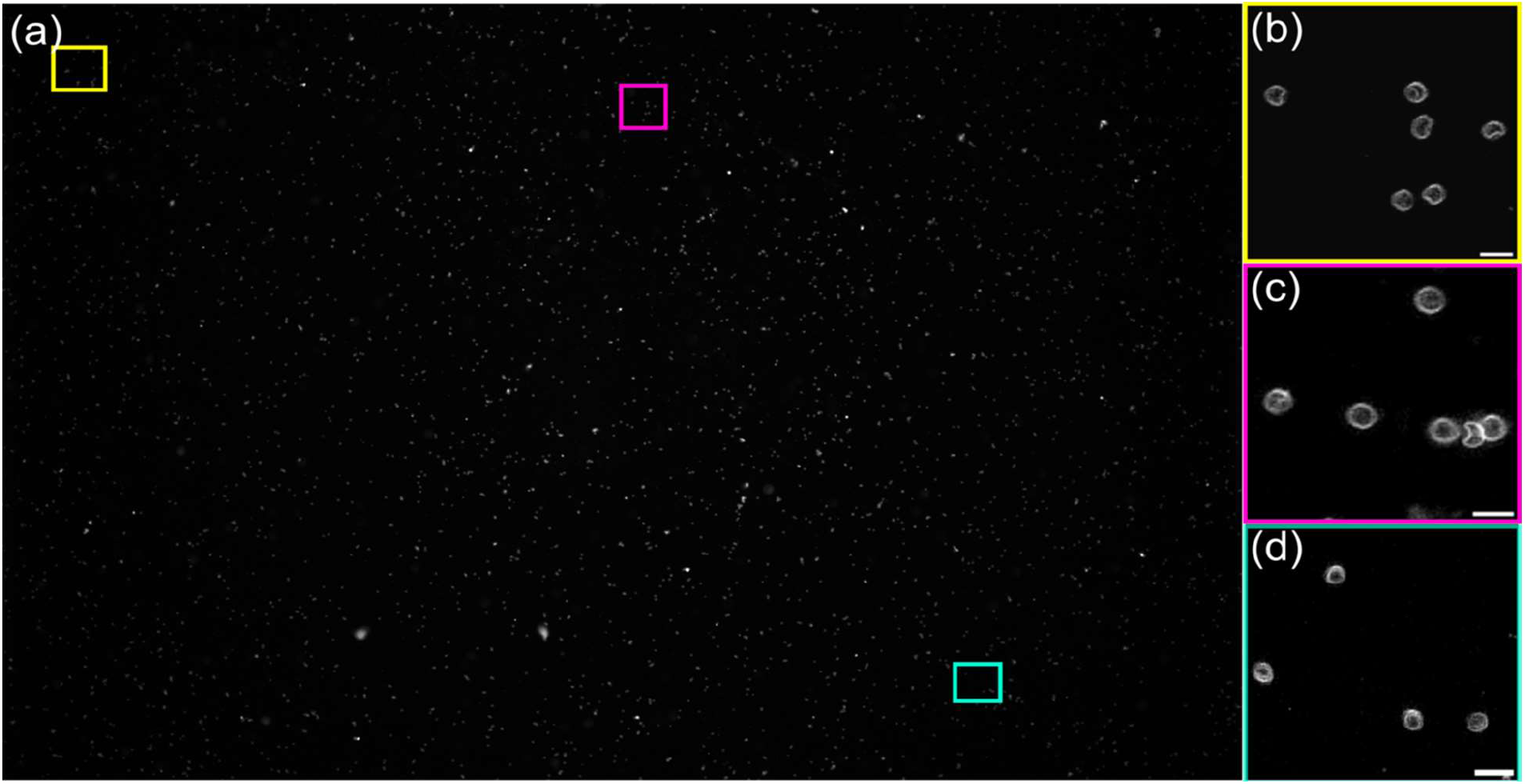
Mesoscale SW image of live red blood cells stained with FM4-64. Three ROIs are shown with a yellow box at the top left of the image, a magenta box close to the centre of the field, and a cyan box towards the bottom right of the field. Scale bar = 1 mm. (b-d) Digital zoom of the ROI shown with the yellow, magenta, and cyan boxes respectively. Scale bars = 10 μm. 4(b)-(d) confirm multi-plane imaging across the large FOV of the Mesolens.

Using the high-volume cell suspension, ten cells (of a population of thousands) moving under flow conditions were cropped from the original data and combined into a single movie. This is shown in Movie 1, in which the playback speed was set to 2 frames per second. Cell movement and topology are clearly visible, with cells travelling at different velocities in different regions of the specimen. For example, at time t=210 s, fluorescence excited by two anti-nodal planes is visible for cells in the first, fourth and fifth rows of the movie. As the time advances, the SW illumination pattern reveals the changing topology of the cells as they rotate in the flowing fluid. No photobleaching is observed in the data, and there are no obvious structural changes to imaged cells, suggesting that any photo-damage to the specimen is negligible.

**Movie 1:**
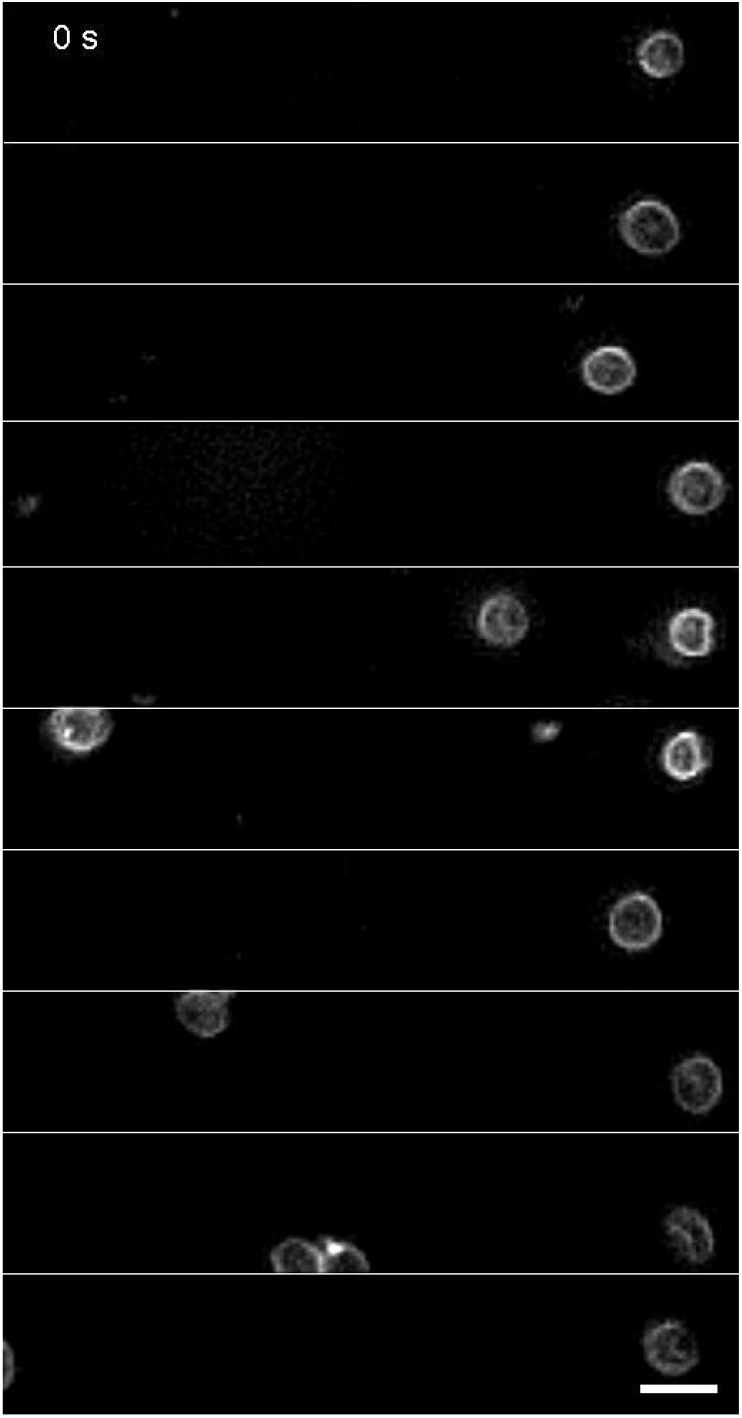
Ten red cells imaged with standing wave microscopy at time t=30 s intervals are cropped from a full FOV SW mesoscopy dataset. The cells are subject to flow conditions, and SW mesoscopy imaging captures cell movement, with cells travelling at different velocities in different regions of the specimen. Scale bar = 10 μm.

## 4. DISCUSSION

The most important findings of our work are that single-wavelength SW imaging and TartanSW imaging can be applied at the mesoscale, and that the Mesolens is suitable for this purpose in spite of the lower NA compared with previously reported SW imaging studies. ^1-3,14^ By preparing specimens using methods previously applied in SW microscopy, we have increased the FOV to 4.4 mm x 3.0 mm, enabling SW imaging of more than 16,000 cells in a single image.

The high numerical aperture of the Mesolens confers high lateral resolution, which means that the topological information from a wide variety of specimens can be easily resolved in the SW image data. However, there are further advantages of the Mesolens over conventional low magnification objectives for SW imaging. For instance, the optical throughput of the Mesolens is 25-times higher than that for a commercial lens with equivalent magnification power. ^6^ Our previous attempts at using commercial low magnification, low numerical aperture lenses for SW imaging with multi-millimetre diameter FOVs failed because the fluorescence signal from the specimens was too low to produce a good quality image, and only images with very low contrast could be obtained. We have shown that the higher optical throughput of the Mesolens, used in combination with the chip-shifting camera, can produce high-contrast, high-quality SW datasets.

The obvious advantage of SW for three-dimensional imaging is the ability to study cell topology in a single image, but a less obvious benefit is the high speed of imaging compared to other three-dimensional imaging methods such as confocal laser scanning microscopy. For instance, for confocal imaging with the Mesolens we note that with a 4.4 mm by 3.0 mm FOV the number of pixels required to fulfil the Nyquist sampling criterion is 19,728 pixels x 13,152 pixels. Using a minimum practical pixel dwell time of ca. 0.5 μs leads to an acquisition time of 130 s for a full FOV, full resolution scanned image, which gives only two-dimensional information from the specimen. Our SW image data, on the other hand, which is also Nyquist sampled across the lateral field and offers insights into the third dimension, is acquired at a rate more than 8 times faster than confocal imaging can provide. This method may facilitate the diagnosis of parasitaemia ^21^ and other diseases of the blood including hereditary spherocytosis, sickle cell disease, hereditary stomatocytosis or elliptocytosis. ^22^

We note that the depth of field of the Mesolens of approximately 8 μm imposes a limitation on the number of anti-nodal planes that can be detected and hence restricts the method to specimens with a maximum thickness of 8 μm.

Our red blood cell SW data is limited to a maximum of two anti-nodal planes that can be resolved in the individual red blood cells. This is contrary to previous work using high magnification, high numerical aperture lenses that showed up to four anti-nodal planes. ^2,14^ We attribute this reduction in anti-nodal planes to the resolving power of the Mesolens: there are likely to be up to four anti-nodal planes illuminating the specimens, but the lateral resolution of the Mesolens is insufficient to distinguish them in the highly curved red cell membrane. We tried to resolve these using different deconvolution methods, but this did not change the overall result.

As with the original TartanSW microscopy method, ^3^ complexities of the mesoscale TartanSW image data preclude straightforward reconstruction of the three-dimensional object structure. Unfortunately, the fluorescently stained actin cytoskeleton proved too complex to be able to reliably resolve the colour difference in anti-nodal planes except in the thinnest part of the cell close to the edge, so resolving adjacent structures in z with a resolution higher than a confocal microscope can provide was not feasible. There are several algorithms for multi-wavelength interferometric surface profiling, but SWs close to a reflector present a much more complex case than the single or multi-wavelength reflections used in surface profilometry. ^23^ As a consequence, these reconstruction algorithms cannot be easily implemented with TartanSW datasets at either the microscale or mesoscale. It would be interesting to modify phase-unwrapping methodologies previously applied to multiwavelength images of protozoa to produce three-dimensional reconstructions of SW images. ^24^

Our time-lapse imaging data raises the possibility of using SW mesoscopy to study cell deformation under lateral flow conditions to understand the role of shear stress on cell health and regeneration. ^12^ To our knowledge, this presents a new application for SW imaging, and the large FOV of the Mesolens would facilitate long-range tracking of large cell populations in three-dimensions within a single dataset. This should ideally be performed in a custom-designed flow chamber that incorporates an in-built first surface reflector.

Although the Mesolens can produce full-resolution images of up to 6 mm in diameter, the FOV is limited to 4.4 mm by 3.0 mm in this study. This is because of limitations of detection, principally the chip-shifting camera. A successor to our chip-shifting camera includes a 50 MP sensor that has a smaller pixel size and is capable of producing images of up to 427 MP. ^25^ Such an improved camera could potentially be applied to extend SW mesoscopy yet further to a FOV up to 6 mm in diameter. These devices use a CMOS chip that can image at up to 3 frames per second, and therefore may further increase both the detection sensitivity and imaging speed compared to the CCD-based device we have used in this work. This may be of considerable benefit for the application of SW mesoscopy to the study of cells in flow conditions.

## 5. CONCLUSIONS

In this study, we have presented single-wavelength SW and multi-wavelength TartanSW imaging at the mesoscale using the Mesolens. The combination of the high optical throughput afforded by the Mesolens, in combination with the low magnification and high numerical aperture, allowed us to perform SW imaging of non-biological specimens and both fixed and live cell specimens. For single-colour SW mesoscopy, two-dimensional datasets that contain three-dimensional information can be obtained in approximately 15 seconds. This is a considerable improvement in acquisition speed compared to other methods such as confocal laser scanning microscopy with the Mesolens or computational stitching and tiling of small data volumes. Our single-wavelength SW and multi-wavelength TartanSW mesoscopy cell images are of high-quality and high-contrast, and we observed no obvious photobleaching or photodamage when imaging live cells. To our knowledge, our application of SW mesoscopy to visualise the membranes of over 16,000 fluorescently-stained red blood cells simultaneously represents the largest cell population that has been observed in a single dataset using SW methods. We show the potential of the method for using SW mesoscopy to study red blood cells under flow conditions, and we discuss how these results may be further improved using new and emerging detector technologies.

## Acknowledgements

This work was supported by the Biotechnology and Biological Sciences Research Council, grant numbers BB/P02565X/1 and BB/T011602/1. S.F. was supported by the Engineering and Physical Sciences Research Council iCASE studentship award, supported by the National Physical Laboratory. L.S.K. was supported by the Medical Research Council and Engineering and Physical Sciences Research Council Centre for Doctoral Training in Optical Medical Imaging, grant number EP/L016559/1. M.S. was supported by the National Measurement System funded by UK’s Department for Business, Energy & Industrial Strategy. G.M. was partly supported by the Medical Research Council, grant number MR/K015583/1 and the Leverhulme Trust.

## Author Contributions

S.F., J.K.S., T.B., and G.M. designed research; S.F. and J.K.S. prepared specimens for imaging; S.F., J.K.S. and G.M. performed SW imaging; S.F., L.S.K., and G.M. performed image processing and analysis; G.M. wrote the article; All authors edited the article.

